# Does offspring sex ratio differ between urban and forest populations of great tits (*Parus major*)?

**DOI:** 10.1101/2020.01.30.921106

**Authors:** Nóra Ágh, Ivett Pipoly, Krisztián Szabó, Ernő Vincze, Veronika Bókony, Gábor Seress, András Liker

## Abstract

Since male and female offspring may have different costs and benefits, parents may use sex ratio adjustment to increase their fitness under different environmental conditions. Urban habitats provide poorer conditions for nestling development in many birds. Therefore, we investigated whether great tits (*Parus major*) produce different brood sex ratios in urban and natural habitats. We determined the sex of nestlings of 126 broods in two urban and two forest habitats between 2012 and 2014 by molecular sexing. We found that brood sex ratio did not differ significantly between urban and forest habitats either at egg-laying or near fledging. Male offspring were larger than females in both habitats. This latter result suggests that male offspring may be more costly to raise than females, yet our findings suggest that urban great tits do not produce more daughters despite the unfavourable breeding conditions. This raises the possibility that other aspects of urban life, such as better post-fledging survival, might favour males and thereby compensate for the extra energetic costs of producing male offspring.

## Introduction

In birds, brood sex ratio is often differ from parity, and the direction and extent of this difference seems to be not random. Females in many birds species appear to optimize the brood sex ratio according to the cost and fitness outcome of producing male and female offspring, which may vary among environments as well as with the quality of the parents (Szász et al. 2012). For example, one sex may have higher growth rate than the other, resulting in sexual size dimorphism (one sex having larger body size than the other). This can be one of the main causes of the unequal costs of male and female offspring to parents (e.g. Martins, 2004; Rosivall et al. 2004; Råberg et al. 2005), as a faster-growing or larger offspring needs larger amounts of food, requiring higher parental effort (e.g. Anderson et al. 1993; Kalmbach et al. 2001). Sexual size dimorphism is widespread in birds, both in eggs (e.g. Cordero et al. 2000, 2001) and in nestlings (e.g. larger females: Anderson et al. 1997; Massemin et al. 2000; larger males: Howe, 1977; Hochachka & Smith, 1991; Badyaev et al. 2001; Tschirren et al. 2003). Sex differences in offspring survival rate also affect their relative values. For example, different sensitivity of the sexes to environmental stressors like parasites may induce higher nestling mortality in one sex compared to the other. The larger sex is more likely to be the more sensitive one, because there may be a trade-off between growth and immunocompetence, and the larger sex may allocate more resources in the former at the expense of the latter (e.g. Tschirren et al. 2003 but see Bize et al. 2005). Furthermore, after fledging, the sexes can greatly differ in their dispersal distance (see examples in Végvári et al. 2018), mortality and lifespan (e.g. Liker & Székely 2005; Barrett & Richardson, 2011). These components of male and female life history can be highly dependent on environmental factors (for theoretical model see Julliard, 2000). Accordingly, the optimal brood sex ratio can differ between different environments. For example, mothers may produce more offspring of the less vulnerable sex in years or habitats with poor dietary conditions, to optimize their parental investment and increase the number of surviving offspring (Korpimäki et al. 2000; Pryke & Rollins, 2012). For instance, Komdeur (1996) found in the Seychelles warbler (*Acrocephalus sechellensis*) that producing more females (which remain longer in their natal territories than males) on low-quality territories reduces the parents’ future breeding success, whereas on high-quality territories female offspring stay as helpers, increasing their parents’ breeding success. Therefore, parents with high-quality territories are more likely to produce daughters whereas on low-quality territories they produce more sons.

Urban and non-urban habitats often differ in quality and structure, leading to cardinal changes in life history and breeding phenology of birds in anthropogenic environments (Hinsley et al. 2008; Chamberlain et al. 2009). For instance, urban birds start breeding earlier and have smaller clutches than those in natural habitats (reviewed in Sepp et al. 2018, examples for great tit: Bailly et al. 2015; Charmantier et al. 2017; Seress et al. 2018). In cities, body condition of fledglings is often lower and their mortality rate is higher, which may be compensated for by better adult survival (reviewed in Chamberlain et al. 2009; Seress & Liker, 2015; Biard et al. 2017). Thus, urbanization may change the relative benefits of male and female offspring, resulting in biased brood sex ratio. In urban environments, reduced availability of natural food sources like arthropods during brood-rearing (see e.g. Seress et al. 2018) may have a stronger negative effect on the faster-growing and larger offspring, making the smaller sex more profitable for parents (for similar effects in non-urbanization context, see Rosivall et al. 2010). Furthermore, competition for arthropod food may continue after fledging and might be stronger in urban habitats with unfavourable local conditions than in forests, which predicts that parental investment should be biased towards the more-dispersing sex (Julliard, 2000). Thus, studying offspring sex ratios may contribute to a better understanding of how animals adapt to urban environments. However, our knowledge regarding sex ratio adjustment in urban environments is still very limited (e.g. Dhondt 1970, Rejt et al. 2005, Bonderud et al. 2017).

Beside environmental conditions, parental quality is another factor that can influence future reproductive success of male and female offspring, and thus may also affect the brood sex ratio. On the one hand, the “mate attractiveness hypothesis” (Burley, 1981, 1986) states that females mating with males with attractive heritable traits should produce more sons than those who mate with unattractive males, because the formers’ sons will be more desirable for females and can achieve higher breeding success (e.g. West et al. 2000; Komdeur & Pen, 2002; Yamaguchi et al. 2004; reviewed in Booksmythe et al. 2017). Larger body size (e.g. as indicated by tarsus length in great tits: Yamaguchi et al. 2004) may be one of these attractive heritable male traits. On the other hand, parents of larger body size or in better condition may provide higher quality parental care, which can also influence parents’ decision on optimal sex allocation. This latter idea predicts that higher-quality parents who can provide adequate care under unfavorable conditions (e.g. can provide more and better prey items to the nestlings) will produce more offspring of the more vulnerable sex than lower-quality parents. This, again, predicts an overproduction of the less sensitive sex in urban broods, because body size, condition, and individual quality is often reduced in urban adults (reviewed in e.g. Seress & Liker, 2015).

In this study, we investigated the effects of urbanization on brood sex ratio in great tits, a passerine bird that occupies a wide range of habitats (Burfield & van Bommel, 2004). Great tits are successful urban colonizers, but in cities they often show reduced clutch size, lower nestling mass and fledging success compared to forest areas (Horak, 1993; Chamberlain et al. 2009; Bailly et al. 2015; Seress et al. 2018), likely because of the lower availability of natural prey as nestling food in urban habitats (Seress et al. 2018). In this species, an earlier study found signs of facultative sex ratio adjustment, as primary sex ratios varied with date and clutch size (Lessells et al. 1996). Other studies suggest that different sensitivity of the sexes to habitat quality can also affect the brood sex ratio in this species. For example, Bouvier et al. (2016) found that the sex ratio of fledglings was more biased towards females in orchards with high levels of pesticide treatments (hence reduced food availability) compared to moderately treated or organic gardens. Similarly, breeding territory quality also may predict brood sex ratio in woodland great tits: Stauss et al. (2005) found that in deciduous forests, where caterpillars (the preferred nestling food) were abundant, broods were more male-biased than in coniferous forests that had reduced caterpillar availability. However, none of the earlier studies investigated habitat-related effects on offspring sex ratios in great tits in an urbanization context. Furthermore, the earlier studies investigated only the fledgling sex ratio (which can be changed by parental adjustment or sex-dependent mortality) and not the primary sex ratio (i.e. sex ratio adjustment by parents).

In great tits male offspring are larger and may be more sensitive to poor environmental conditions (Tschirren et al. 2003), whereas females disperse further and thereby may escape more successfully from unfavourable local conditions (Andreu & Barba, 2006). So based on the aforementioned results, we predicted that great tits would produce more female-biased broods in the food-limited urban habitats than in natural forests where nestling food is abundant. We tested this prediction using breeding data from three years of monitoring four populations, two in cities and two in nearby deciduous woodlands. We investigated both the primary sex ratio (i.e. sex ratio at egg laying) and fledgling sex ratio, and we took into account other factors that may influence brood sex ratios, including laying date and, as proxy for parental quality, parental body size (e.g. Kölliker et al. 1999; Rosivall et al. 2004; Bell et al. 2014). Using data on fledgling body size and nestling survival, we also evaluated whether male offspring are larger and more sensitive (in terms of nestling mortality) than females in our populations.

## Materials and Methods

### Field methods

We studied great tit populations at two forests and two urban sites in Hungary. Forest sites were located in deciduous woodlands near Szentgál (47°06’39.75”N, 17°41’17.94”E) and in Vilma-puszta (47°05’06.7”N, 17°51’51.4”E), whereas the two urban sites were located in the cities of Veszprém (47°05’17.29”N, 17°54’29.66”E) and Balatonfüred (46°57’30.82”N, 17°53’34.47”E). We collected data at all study sites from 2012 to 2014, with the exception of Balatonfüred, where data collection started in 2013. Nest boxes in the urban habitats were placed mostly in public parks and university campuses; all of these plots were strongly influenced by anthropogenic disturbance (e.g. presence of vehicle traffic and human activitiy; see Seress et al 2018 for more details on the study sites). We monitored the nest boxes at least twice a week from March to early July to record laying date of the first egg, clutch size, hatching dates, and the number of nestlings (detailed in Seress et al. 2017). We ringed all nestlings just before fledging (at 14-16 days of age, day 1 being the hatching day of the first-hatching nestlings) and measured the length of their left tarsus to the nearest 0.1 mm and their right wing (the flattened maximum wing chord, from the carpus to the tip of the longest primary; Svensson, 1992) to the nearest mm, and recorded their body mass (to the nearest 0.1 g using Pesola spring balance). We also took a small drop of blood (ca. 25 μl) from the brachial vein. In 2013-2014, we collected unhatched eggs (that did not hatch for at least 5 days after the first chick of the same brood hatched) and a small tissue sample (e.g. feather, toes) from chicks found dead in the nest during nest box checking throughout the brood rearing period. We stored all samples either in Queen’s lysis solution or in 96% ethanol at 4°C until further analysis. We captured adult birds on their nests during brood rearing and ringed each bird with a unique combination of a numbered metal ring and three plastic colour rings for individual identification (Seress et al. 2017). To increase the number of individually identified birds in our populations, we also ringed adult great tits outside of the breeding season (from late September to early February) at the four study sites using mist-nets. Thus, parents of the broods included in our analyses were identified either by capturing them during brood rearing or by observing their colour ring combinations from video recordings filmed with concealed nest cameras (see Seress et al. 2017 for details). On these video samples we considered a colour-ringed individual to be a parent bird if it was recorded to enter the nest box with food at least once. For measuring and sampling adult birds, we followed the same protocol described above for fledglings.

### Laboratory methods

We extracted DNA by using silica membrane isolation kits (GeneJET, Genomic DNA Purification Kit) following the manufacturers’ protocol (Thermo Scientific™). Molecular sexing was performed using the primer pairs P2 – P8 with the protocol of Griffiths et al. (1998). We investigated all unhatched eggs for the presence of an embryo before DNA isolation. If we noticed no sign of embryo development (not even a visible germinal disc), we classified them as infertile eggs. Out of 44 unhatched eggs, we found 30 infertile eggs. We preserved the embryos from the 14 fertile eggs in 96% ethanol. We then extracted a small sample of tissue from the embryos and the further DNA isolation steps were similar to the methods we used for blood and other tissue samples. All embryos were successfully sexed. We were also able to successfully extract DNA from all of the tissue samples of the dead nestlings.

We analysed 126 broods (14 from 2012, 52 from 2013, and 60 from 2014) where we had blood or other tissue samples from nearly all offspring (i.e. missing tissue sample from no more than 3 dead offspring per brood). We had 79 broods (6 from 2012, 34 from 2013, and 39 from 2014) where we were able to take DNA samples from all offspring (both dead and fledged) and thereby we could calculate the primary sex ratio (i.e. at egg laying). The 6 broods from 2012 that we could include in the primary sex ratio analyses were nests where all laid eggs had become successful fledglings (i.e. there were no unhatched eggs or dead nestlings). In the remaining broods we could estimate only the fledgling sex ratio (i.e. at the age of ringing, at 14-16 days). We aimed to sample both the first and second annual broods at each study site. We categorized each brood as the first annual breeding attempt of a pair if it was initiated before the date of the first egg laid in the earliest identified second clutch in that year at that study site (i.e. clutch by a colour-ringed female that had already successfully fledged at least one young in that year). Broods initiated after this date were categorized as second annual breeding attempts. Our sample size is inherently unbalanced, because the number of available broods differed between sites and years, and changed over the season (i.e. there were fewer second broods than first broods). For the 126 broods, we were able to identify 240 parents, out of which 111 fathers and 118 mothers were measured as adults (the remaining 11 birds were only measured and ringed as nestlings in the previous year); in total, we had 105 broods were both parents were identified and measured.

### Statistical analyses

We calculated primary and fledgling sex ratios as number of males divided by the total number of offspring/nestlings. Primary sex ratio means the sex ratio of all offspring (embryos, dead chicks, and chicks that reached the fledging age) in complete broods, whereas fledgling sex ratio means the sex ratio of nestlings that reached the fledging age (without embryos or dead chicks). We analysed the data from the first and second annual broods together and used the laying date as a covariate in all analyses. We calculated laying date in two alternative ways, and used these two variables in two alternative sets of models. First, we used laying date as the absolute number of days since 1 January until the laying of the first egg in the brood (Julian day). This variable reflects brood value, as offspring fledging later in the season have less time for post-fledging growth before winter. Second, to test the specific effect of timing within the breeding season in each year at each study site, we used mean-centered laying date, subtracting the mean of the respective site and year from each brood’s laying date. This variable captures a different aspect of the date effect, as the start of the breeding season varies among sites and years, and the relative timing of broods may affect their food availability (Seress et al. 2018). In the main text, we present the results using the former date variable; see the Supplementary Material for results with the latter date variable (Table S3).

To test whether the primary and fledgling sex ratios differed between study sites, we built generalized linear mixed-effects models with binomial error distribution and “logit” link function (function glmmPQL in package MASS; Ripley et al. 2013). The full models contained study site, year, laying date (either Julian day or the mean-centered laying date), tarsus length of the father, and tarsus length of the mother as fixed effects and brood ID nested in pair ID as random factors. We also tested the interaction between study site and parents’ tarsus length, but it was non-significant in all models (*P* > 0.08), so we present all model results without these interactions to facilitate easier interpretation of the main effects. Note that we did not include other parental body size variables (i.e. wing length, body mass) as predictors of brood sex ratio, because these traits can change considerably throughout the year and in many cases parents’ size data were collected outside of their breeding period (see Field methods above). To increase our sample size, we repeated these analyses after excluding parents’ tarsus length from the model, because we had data on both parents’ tarsus length only in a subset of broods (see Supplementary Table S1 for sample sizes). Henceforward we refer to these analyses as “reduced models”. Furthermore, to assess if our results were affected by imbalanced sample sizes due to the different frequency of second annual broods at the four sites, we repeated our sex ratio analyses after excluding the second broods.

To statistically compare the sex ratios between the two habitat types (urban sites vs. forest sites) we calculated linear contrasts from the full and reduced models. These linear contrasts were pre-planned comparisons between the two urban sites vs. the two forest sites (see also Pipoly et al. 2019 and Vincze et al. 2019 for the same approach to compare habitat types by pre-planned linear contrasts and for additional details of the method). Each linear contrast was back-transformed from the log-scale to provide the odds ratio (OR, i.e. the proportional difference of the odds of an offspring being male between urban and forest broods) with 95% confidence interval (CI). For the linear contrasts, we used the “emmeans” function (emmeans package in R; Lenth & Lenth, 2018).

To investigate sexual size dimorphism in fledglings (measured at ringing, 14-16 days post-hatching; day of hatching = day 1), we used linear mixed-effects models (function lmer in package lme4; Bates et al. 2014). We built three separate models in which the response variables were the wing length, tarsus length or body mass of individual fledglings, respectively. In these three models the fixed effects were study site, year, laying date (Julian day only) and sex of the fledgling, while brood ID nested in pair ID and crossed with measurer ID were included as random factors. To test if body size differences between male and female fledglings were different at the four study sites, we added the two-way interaction between sex and study site to these models. Any random variation among broods (including any difference in age) was taken into account by including brood ID as a random factor. We did not include fledgling age at ringing into the model because it varied in a very narrow range (14-16 days); note that Seress et al. (2018) found no significant effect of fledgling age (within the same age interval) on body mass in the same populations between 2013 and 2016.

To test for sex-dependent offspring survival, we analysed the effect of offspring sex on the probability of mortality to fledging. We used a generalized linear mixed-effects model with binomial error distribution and “logit” link function glmmPQL in package MASS (Ripley et al. 2013). The response variable was the status of offspring as alive (survived to day 14-16) or dead (unhatched eggs and dead chicks), the fixed effect was the sex of the offspring, and the model also included brood ID nested within pair ID as random factors. Because offspring mortality was very rare, especially in forest sites (see Results), we did not investigate whether the sex difference in mortality differed between habitats.

All of the tested variables showed acceptable level of multicollinearity, because the variance inflation factor (VIF) varied from 1.04 to 1.22 in all of the models. All analyses were done using R version 3.4.2. (R Core Team 2017).

## Results

In our sample, primary sex ratio was overall 0.493, whereas fledgling sex ratio was 0.514 (for sample sizes see Table S1). For both primary and fledgling sex ratio, none of the tested predictors had significant effects either in the full model (see model estimates in Table S2 & S3) or in the reduced model (Table 1). Primary sex ratio was statistically close to parity at every study site (estimated mean ± SE, Veszprém city: 0.55±0.248; Balatonfüred city: 0.46±0.390; Vilma-puszta forest: 0.46±0.256; Szentgál forest: 0.48±0.246; the 95% CI includes 0.5 for all sites, see Figure 1) and did not differ significantly between urban and forest sites (Table 1). Fledgling sex ratio also did not deviate significantly from parity at any of the four sites (Veszprém city: 0.60 ±0.171; Balatonfüred city: 0.51±0.483; Vilma-puszta forest: 0.51±0.265; Szentgál forest: 0.52±0.248; the 95% CI includes 0.5 for all sites, see Figure 2), and there was no significant difference between urban and forest habitats (Table 2). These results were qualitatively identical when we eliminated the second annual broods from the models (see model estimates in Table S4 and Table S5).

**Table 1:** Primary and fledgling sex ratio of great tits in relation to study site, year, and laying date (Julian day, first and second annual broods pooled). Effects are presented as analysis of deviance tables with type-2 sums of squares for the reduced generalized mixed-effects models; n= 79 and 126 for primary and fledgling sex ratios, respectively.

| | $\chi^2$ | df | *P* |
| --- | --- | --- | --- |
| **Primary sex ratio** |  |  |  |
| Sites | 2.040 | 3 | 0.564 |
| Years | 0.036 | 2 | 0.982 |
| Laying date | 1.655 | 1 | 0.198 |
| **Fledgling sex ratio** |  |  |  |
| Sites | 1.707 | 3 | 0.635 |
| Years | 0.430 | 2 | 0.807 |
| Laying date | 2.563 | 1 | 0.109 |

**Figure 1:**
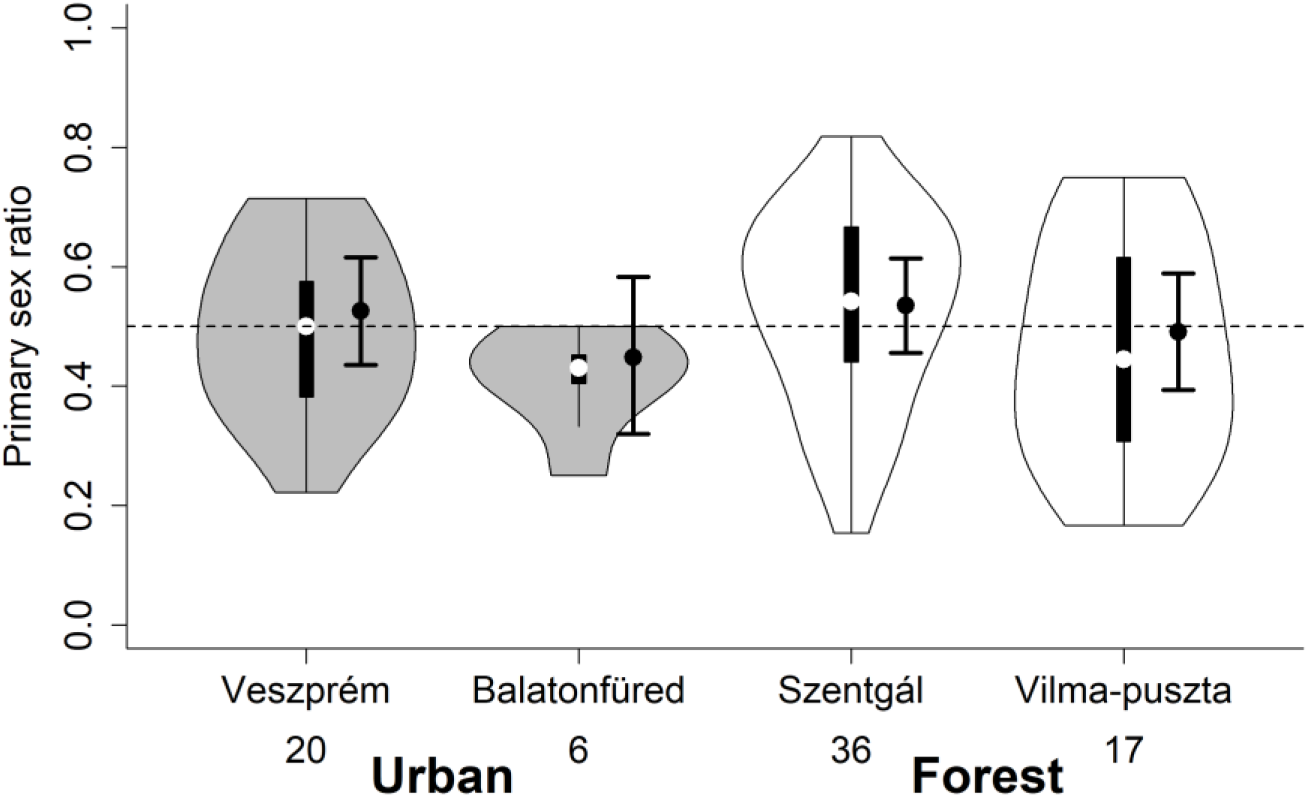
Violin plot of the distribution of primary sex ratio (proportion of males) in broods at urban and forest study sites (first and second annual broods pooled). Each plots show the median (indicated by the small, open circle), the first through the third interquartile range (the thick, solid vertical band), and estimator of the density (thin vertical curves) at each site. Numbers below the violin plots refer to the number of broods in each site. Dots and whiskers next to the inner box plots show means and 95% confidence intervals, respectively, both calculated from the model shown in Table 1.

**Table 2:** Differences (pre-planned linear contrasts) in primary and fledging sex ratios between urban and forest habitats, shown as odds ratio (OR; proportional difference of the odds of an offspring being a male at urban sites compared to forests).

|  | OR [95%CI] | df | t | *P* |
| --- | --- | --- | --- | --- |
| **Primary sex ratio** |  |  |  |  |
| Full model | 0.87 [0.71;1.06] | 53 | -0.697 | 0.489 |
| Reduced model | 0.90 [0.75;1.09] | 72 | -0.549 | 0.584 |
| **Fledgling sex ratio** |  |  |  |  |
| Full model | 1.04 [0.88;1.23] | 96 | 0.236 | 0.814 |
| Reduced model | 1.07 [0.92;1.25] | 104 | 0.473 | 0.637 |

**Figure 2:**
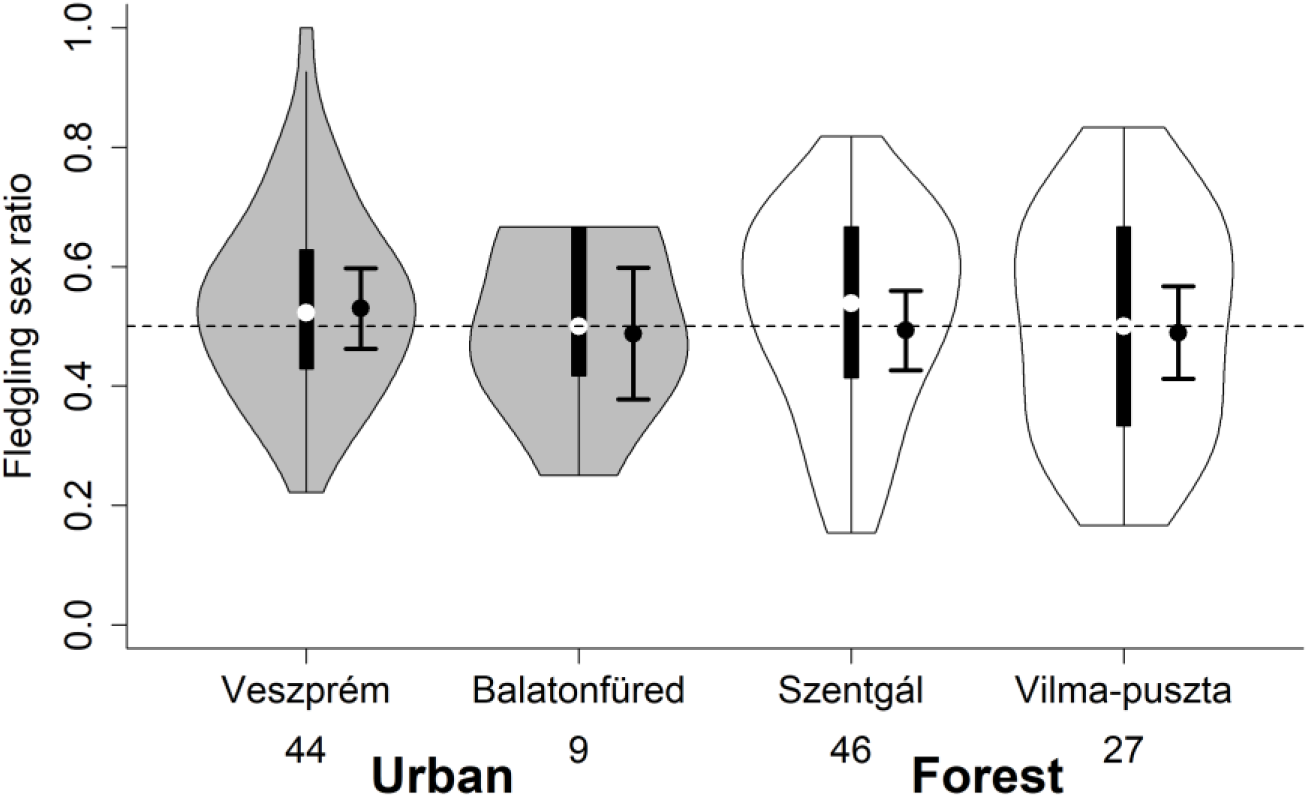
Violin plot of the distribution of fledgling sex ratio (proportion of males) in broods at urban and forest study sites (first and second annual broods pooled). Each plots show the median (indicated by the small, open circle), the first through the third interquartile range (the thick, solid vertical band), and estimator of the density (thin vertical curves) at each site. Numbers below the violin plots refer to the number of broods in each site. Dots and whiskers next to the inner box plots show means and 95% confidence intervals, respectively, both calculated from the model shown in Table 1.

Male fledglings had longer wings and tarsi and were heavier than female fledglings (Figure 3, Table 3). These size differences between sexes were independent from the study site (interactions between the sex of the nestlings and study site were non-significant, Table 3). None of the body size parameters varied significantly with laying date or among years (Table S6).

**Table 3:**
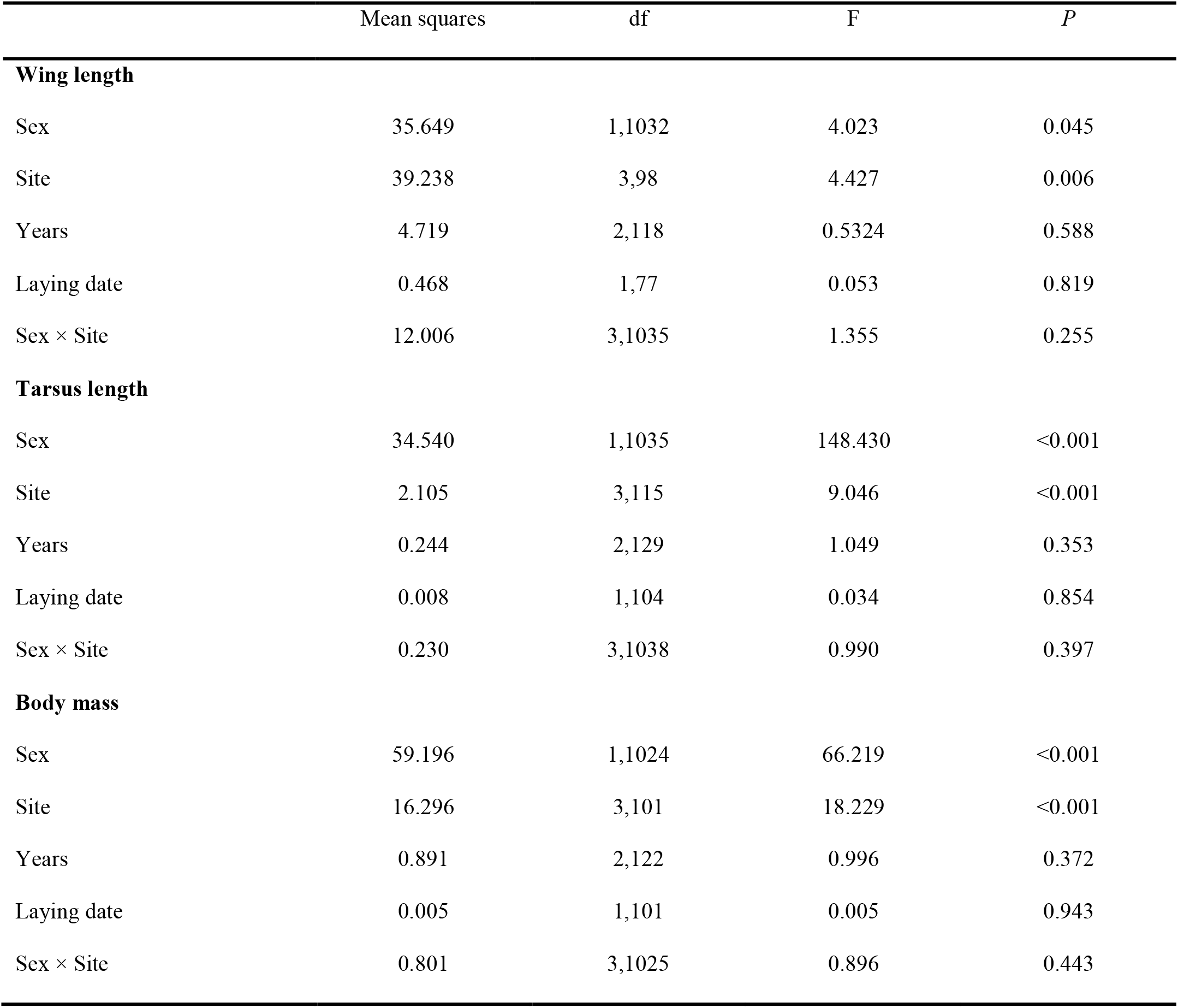
Results of the analyses of body size parameters of nestlings in relation to their sex, study site, years and laying date. Effects are presented as analysis of deviance tables with type-2 sums of squares for the reduced generalized mixed-effects models. Nestlings of first (n= 952) and secoond annual broods (n= 200) were pooled in the analyses.

|  | Mean squares | df | F | *P* |
| --- | --- | --- | --- | --- |
| **Wing length** |  |  |  |  |
| Sex | 35.649 | 1,1032 | 4.023 | 0.045 |
| Site | 39.238 | 3,98 | 4.427 | 0.006 |
| Years | 4.719 | 2,118 | 0.5324 | 0.588 |
| Laying date | 0.468 | 1,77 | 0.053 | 0.819 |
| Sex × Site | 12.006 | 3,1035 | 1.355 | 0.255 |
| **Tarsus length** |  |  |  |  |
| Sex | 34.540 | 1,1035 | 148.430 | <0.001 |
| Site | 2.105 | 3,115 | 9.046 | <0.001 |
| Years | 0.244 | 2,129 | 1.049 | 0.353 |
| Laying date | 0.008 | 1,104 | 0.034 | 0.854 |
| Sex × Site | 0.230 | 3,1038 | 0.990 | 0.397 |
| **Body mass** |  |  |  |  |
| Sex | 59.196 | 1,1024 | 66.219 | <0.001 |
| Site | 16.296 | 3,101 | 18.229 | <0.001 |
| Years | 0.891 | 2,122 | 0.996 | 0.372 |
| Laying date | 0.005 | 1,101 | 0.005 | 0.943 |
| Sex × Site | 0.801 | 3,1025 | 0.896 | 0.443 |

**Figure 3:**
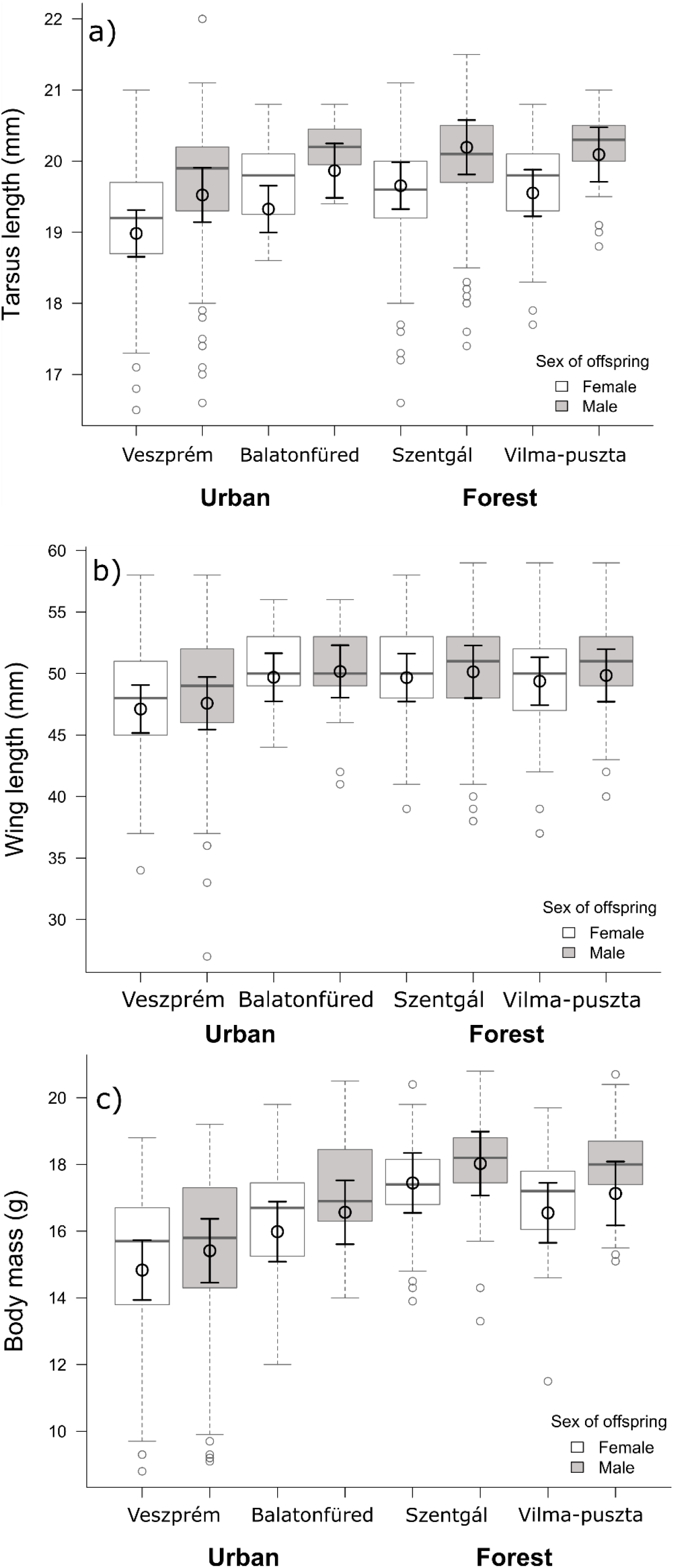
Body size (a: tarsus length, b: wing length, and c: body mass) of male and female fledglings at the study sites. Box plots show the median, lower and upper quartiles and the whiskers represent data within the 1.5 × interquartile range. The error bars show the mean ± SE values estimated from the linear mixed models in Table 3. Details on parameter estimates for sex and site effects are provided in Table S6.

In our sample, 10 males and 4 females from 10 broods died in the egg, and 7 male and 6 female nestlings from 9 broods died before ringing. The highest number of dead offspring was found in Veszprém (n= 17), whereas at the other sites mortality was very low (Balatonfüred: n= 5, Szentgál: n= 3, Vilma-puszta: n= 2). The sex ratio of dead offspring was 0.63 (0.59 in cities and 0.80 in forests); the proportional difference of the odds of mortality did not differ significantly between males and females (OR= 1.50, CI= 0.91 – 2.47, *P*= 0.411).

## Discussion

Contrary to our prediction that great tit parents may overproduce daughters in food-limited urban habitats, we found that neither the primary nor the fledgling sex ratios differed signifiantly between urban and forest study sites. We consider these results robust, because we collected data over three breeding seasons at four study sites (two urban, two forest), and excluding the second annual broods did not change our results qualitatively (Tables S4 & S5). Our results differ from the findings of two other studies comparing great tits’ offspring sex ratios between habitats of different quality. In one of these earlier studies, where the sexing of nestlings was based on visual cues (Dhondt 1970), more male offspring were found in urban compared to suburban or woodland habitat before fledging. In the other study, Bouvier et al. (2016) found more male nestlings in organic orchards with less pesticide use (that likely represent better habitat quality) than in orchards cultivated by using large amounts of pesticide. The reason for the varying results among these studies is unclear. Notably, the aforementioned studies showed information only about fledgling sex ratio, so to our knowledge our study is the first that compare primary sex ratio between urban and forest habitats in great tits.

With the available information, we can only speculate why we did not find sex ratio adjustment in urban habitats. First, it is possible that in our study populations male or female offspring did not differ in the associated costs of producing and raising them until independence. However, 14-16 days old male fledglings were significantly heavier (by 3.6%) and had slightly longer tarsi (by 2.5%) and wings (by 2%) compared to their female siblings, regardless of habitat type. These results suggest that male nestlings require more parental provisioning during their development than females, although we do not know the extent (and hence the associated additional costs) of such extra provisioning. Apparently, parents were able to meet this requirement in both habitats, because the size difference between male and female fledglings was similar in all study sites, and we did not find any evidence for sex-related mortality. This seems to contradict earlier studies in other great tit populations, which reported either male-biased sex ratio in unhatched eggs (Cichoń et al. 2005) or higher mortality in females before fledging (e.g. Smith et al. 1989; Lessells et al. 1996), and in some cases growth of females was more severely affected by poor condition in tit species (Oddie 2000, Nomi et al. 2018). To better understand these conflicting results, we need to have more data on the sex-specific mortality rates before and after hatching from our study populations and also on the environmental factors and parental quality variables that can influence embryo and nestling survival. For example, it is possible that the increased resource requirement of male offspring induces male-biased mortality only under unusually poor conditions, such as harsh weather, high prevalence of parasites or disease, or extremely low food supply (Tschirren et al. 2003).

Given that the larger size of male fledglings suggests higher parental cost, a potential explanation for the lack of sex ratio adjustment is that there may be some unknown cost to producing female offspring that cancels out the differences in the pay-off between the sexes. For example, it is possible that survival chances are lower after fledging for females than for males. The most dangerous period in the life of juvenile great tits is the dispersion after fledging: Naef-Daenzer et al. (2001) found that 47% of the juveniles died during the first 20 days after fledging. Female great tits disperse farther than males (Andreu & Barba, 2006), which may mean higher risk of mortality for females, especially in urban habitats where the potential breeding and feeding sites are more fragmented by built-up areas and roads with heavy traffic. Furthermore, survival during autumn and winter may also differ between the sexes in a habitat-dependent manner. In urban areas, seeds and other food in artificial feeders can increase the chance of survival (Marzluff, 2017), but competition at these feeders can be stronger than at natural feeding sites such as tree canopies. At these feeders, social rank can limit access to food, because subordinate individuals may be attacked by dominant ones and therefore get less food. In great tits, males are more often dominant than females, especially in juveniles (e.g. Barluenga et al. 2000; Dingemanse & de Goede, 2004). These sex differences in great tit life history may generate female-biased mortality, especially in urban habitats. However, the only published study that compared the sex-specific survival of great tits in both urban and rural habitats found higher adult female than male survival in both habitats, and yearling females outnumbered yearling males in next year in breeding season (Hõrak & Lebreton, 1998).

We found remarkably high variance of sex ratios among individual broods in both habitat types (primary sex ratio, range in urban habitat: 0.22 – 0.71, in forest habitat: 0.15 – 0.82; fledgling sex ratio, urban: 0.22 – 1.00, forest: 0.15 – 0.84). This variance in our data was not explained by laying date and the parents’ tarsus length, representing proxies for seasonal environmental changes and for parental quality, respectively. One interpretation of this high variance is that parents vary in their investment into their offspring’s sex, but their allocation is determined by factors which we did not investigate. For example, Lessels et al. (1996) reported that the proportion of male offspring increased with hatching asynchrony in great tits. Furthermore, Pipoly et al. (2019) found in the same populations and breeding seasons as in the present study that the number of extra-pair offspring was higher in urban habitats than in forests, which might influence sex ratio adjustment. The other possible interpretation of our findings is that the observed variance in brood sex ratios is largely random, with no facultative sex ratio adjustment going on (Ewen et al. 2004). For example, in urban areas, where the environmental changes may be rapid and unpredictable, sex ratio manipulation might not be a profitable strategy, as it may be difficult for parents to predict the conditions their offspring will find themselves in. So far, there have been very few studies on great tit primary sex ratios, and their results provided little if any evidence that the observed variation among nests is adaptive (Lessels et al. 1996; Kabasakal & Albayrak, 2012).

## Conclusion for Future Biology

Taking our results together with the small number of previous findings, the role of facultative sex ratio adjustment in birds’ adaptation to urban life is not yet clear. Further studies are needed to better understand the prevalence and drivers of offspring sex ratio in an urbanization context. For example, we should study different environmental predictors that may differ between antropogenic and natural habitats and may lead to differences in the costs and benefits of male and female offspring, influencing sex ratio adjustment. Research is needed also on the sex-dependent effects of urbanization on life-history traits and thus the fitness pay-offs of producing sons and daughters along the urbanization gradient, including sex-related post-fledging survival and future breeding success of male and female offspring in different habitat types. Furthermore, urbanization may interact with other anthropogenic influences such as climate change, potentially resulting in complex effects on sex ratios if males and females differ in their sensitivities to these various perils.

## Acknowledgments

The authors hereby thank all former and current members of the MTA-PE Evolutionary Ecology Research Group who contributed to the fieldwork producing the data for this study. We are grateful to the members of Conservation Genetics Research Group of the University of Veterinary Medicine who helped in the laboratory work.

## Ethical Statement

All applicable international, national and/or institutional guidelines for the care and use of animals were followed. Research was permitted by the Middle Transdanubian Inspectorate for Environmental Protection, Natural Protection and Water Management (permission number: 31559/ 2011). All procedures were in accordance with the guidelines for animal care outlined by ASAB/ABS and Hungarian laws.

## Authors’ Contributions

IP, EV, VB, GS and AL collected the data in the field. NÁ, IP and KSZ conducted the molecular work. NÁ, IP, EV, VB, GS and AL participated in conceptualization, design and data analysis. All authors wrote the manuscript.

## Supplementary Material

**Supplementary Table S1:** Number of broods and offspring used in different models. Full models include parents’ tarsus length as covariate, whereas tarsus length was excluded from reduced models.

| Sex ratio | Model | Broods | Offspring |
| --- | --- | --- | --- |
| **First and second annual broods together** |  |  |  |
| Primary | Full | 62 | 622 |
|  | Reduced | 79 | 793 |
| Fledgling | Full | 105 | 943 |
|  | Reduced | 126 | 1153 |
| **First annual broods only** |  |  |  |
| Primary | Full | 46 | 498 |
|  | Reduced | 59 | 642 |
| Fledgling | Full | 81 | 772 |
|  | Reduced | 98 | 956 |

**Supplementary Table S2:** Primary sex ratio (n= 62 broods) and fledgling sex ratio (n= 105 broods) in relation to site, year, laying date and parents’ tarsus length (first and second annual broods pooled). Estimates with SE were calculated from the parameter estimates of generalized linear mixed-effects models with binomial error distribution and “logit” link function, with brood ID nested in pair ID as random factors. Year parameters show the difference from 2012.

| Model parameters | Estimate±SE | t | P |
| --- | --- | --- | --- |
| **Primary sex ratio** |  |  |  |
| site: Veszprém city | 2.21±3.881 | 0.570 | 0.5713 |
| site: Balatonfüred (city) | 1.96±0.318 | 0.491 | 0.4321 |
| site: Vilma-puszta (forest) | 2.00±0.264 | 0.504 | 0.4325 |
| site: Szentgál (forest) | 2.45±0.246 | 0.612 | 0.3328 |
| year: 2013 | 0.13±0.338 | 0.390 | 0.7103 |
| year: 2014 | 0.32±0.324 | 0.987 | 0.3795 |
| Laying date (Julian day) | 0.006±0.005 | 1.229 | 0.2865 |
| Mother's tarsus length | -0.100±0.144 | -0.701 | 0.4866 |
| Father's tarsus length | -0.058±0.163 | -0.359 | 0.7378 |
| **Fledgling sex ratio** |  |  |  |
| site: Veszprém city | -0.48±3.243 | -0.148 | 0.8824 |
| site: Balatonfüred (city) | -0.69±0.257 | -0.209 | 0.8350 |
| site: Vilma-puszta (forest) | -0.70±0.198 | -0.209 | 0.8349 |
| site: Szentgál (forest) | -0.56±0.190 | -0.167 | 0.8681 |
| year: 2013 | 0.120±0.234 | 0.513 | 0.6169 |
| year: 2014 | 0.101±0.222 | 0.455 | 0.6564 |
| Laying date (Julian day) | 0.004±0.004 | 1.081 | 0.2994 |
| Mother's tarsus length | -0.041±0.116 | -0.352 | 0.7256 |
| Father's tarsus length | 0.045±0.128 | 0.352 | 0.7307 |

**Supplementary Table S3:** Primary and fledgling sex ratio of great tits in relation to study site, year, and laying date mean-centered for study site and year (first and second annual broods pooled). Effects are presented as analysis of deviance tables with type-2 sums of squares for the reduced generalized mixed-effects models; n= 79 and 126 for primary and fledgling sex ratios, respectively.

| | $\chi^2$ | df | *P* |
| --- | --- | --- | --- |
| **Primary sex ratio** |  |  |  |
| Sites | 3.354 | 3 | 0.340 |
| Years | 0.270 | 2 | 0.874 |
| Mean-centered laying date | 1.583 | 1 | 0.208 |
| **Fledgling sex ratio** |  |  |  |
| Sites | 0.994 | 3 | 0.803 |
| Years | 3.591 | 2 | 0.166 |
| Mean-centered laying date | 2.425 | 1 | 0.119 |

**Supplementary Table S4:** Primary sex ratio (n= 59 broods) and fledgling sex ratio (n= 98 broods) in relation to site, year, and laying date (first annual broods only). Estimates with SE were calculated from the parameter estimates of generalized linear mixed-effects models with binomial error distribution and “logit” link function, with brood ID nested in pair ID as random factors. Year parameters show the difference from 2012.

| Model parameters | Estimate±SE | t | P |
| --- | --- | --- | --- |
| **Primary sex ratio** |  |  |  |
| site: Veszprém (city) | -1.64±2.443 | -0.672 | 0.5042 |
| site: Balatonfüred (city) | -2.04±2.461 | -0.830 | 0.4104 |
| site: Vilma-puszta (forest) | -1.92±2.569 | -0.749 | 0.4574 |
| site: Szentgál (forest) | -1.72±2.548 | -0.677 | 0.5014 |
| year: 2013 | -0.07±0.364 | -0.180 | 0.8582 |
| year: 2014 | 0.26±0.520 | 0.492 | 0.6249 |
| Laying date (Julian day) | 0.02±0.024 | 0.682 | 0.4982 |
| **Fledgling sex ratio** |  |  |  |
| site: Veszprém city | -1.75±1.380 | -1.266 | 0.2087 |
| site: Balatonfüred (city) | -2.01±1.433 | -1.399 | 0.1650 |
| site: Vilma-puszta (forest) | -2.043±1.449 | -1.410 | 0.1619 |
| site: Szentgál (forest) | -2.08±1.465 | -1.421 | 0.1588 |
| year: 2013 | -0.06±0.259 | -0.231 | 0.8387 |
| year: 2014 | 0.225±0.325 | 0.693 | 0.5600 |
| Laying date (Julian day) | 0.020±0.014 | 1.414 | 0.2929 |

**Supplementary Table S5:** Urban-forest differences (linear contrasts) in sex ratio of first annual broods. OR refers to the odds ratio of an offspring being male instead of female at urban sites opposed to forest sites. Full models contain the parents’ tarsus length as covariate.

| Urban vs. forest | OR [95%CI] | df | t | P |
| --- | --- | --- | --- | --- |
| **Primary sex ratio** |  |  |  |  |
| Full model | 0.95 [0.73;1.24] | 37 | -0.178 | 0.860 |
| Reduced model | 0.98 [0.78;1.24] | 52 | -0.079 | 0.937 |
| **Fledgling sex ratio** |  |  |  |  |
| Full model | 1.17 [0.97;1.42] | 72 | 0.859 | 0.393 |
| Reduced model | 1.20 [1.00;1.43] | 90 | 1.055 | 0.294 |

**Table S6:** Body size parameters of nestlings in relation to sex, site, year, and laying date (Julian day; first and second annual broods pooled). Parameter estimates with standard error (SE) are shown from linear mixed-effects models, with brood ID nested in pair ID and crossed with measurer ID as random factors. Site parameters show the differences from Veszprém, year parameters show the difference from 2012; the intercept refers to Veszprém 2012.

| Model parameters | Estimate±SE | t | P |
| --- | --- | --- | --- |
| **Wing length (mm)** |  |  |  |
| Intercept | 47.11±1.951 | 24.149 | <0.001 |
| sex: male | 0.47±0.18 | 2.546 | 0.011 |
| site: Balatonfüred (city) | 2.58±1.30 | 1.992 | 0.049 |
| site: Vilma-puszta (forest) | 2.27±0.87 | 2.603 | 0.002 |
| site: Szentgál (forest) | 2.56±0.83 | 3.098 | 0.011 |
| year: 2013 | 0.44±1.11 | 0.397 | 0.692 |
| year: 2014 | 0.98±1.08 | 0.910 | 0.365 |
| Laying date (Julian day) | -0.004±0.017 | -0.259 | 0.796 |
| **Tarsus length (mm)** |  |  |  |
| Intercept | 19.03±0.327 | 58.224 | <0.001 |
| sex: male | 0.47±0.030 | 15.509 | <0.001 |
| site: Balatonfüred (city) | 0.27±0.210 | 1.307 | 0.194 |
| site: Vilma-puszta (forest) | 0.52±0.139 | 3.731 | <0.001 |
| site: Szentgál (forest) | 0.61±0.134 | 4.548 | <0.001 |
| year: 2013 | 0.11±0.178 | 0.627 | 0.532 |
| year: 2014 | 0.23±0.175 | 1.315 | 0.191 |
| Laying date (Julian day) | -0.001±0.003 | -0.214 | 0.831 |
| **Body mass (g)** |  |  |  |
| Intercept | 14.83±0.898 | 16.512 | <0.001 |
| sex: male | 0.58±0.059 | 9.856 | <0.001 |
| site: Balatonfüred (city) | 1.15±0.589 | 1.959 | 0.053 |
| site: Vilma-puszta (forest) | 1.72±0.389 | 4.410 | <0.001 |
| site: Szentgál (forest) | 2.61±0.370 | 7.064 | <0.001 |
| year: 2013 | 0.04±0.506 | 0.087 | 0.931 |
| year: 2014 | 0.47±0.483 | 0.972 | 0.333 |
| Laying date (Julian day) | -0.001±0.008 | -0.082 | 0.935 |

